# A high-throughput amplicon sequencing approach for population-wide species diversity and composition survey

**DOI:** 10.1101/2020.10.12.336545

**Authors:** WT Tay, LN Court, S Macfadyen, F Jacomb, S Vyskočilová, J Colvin, PJ De Barro

## Abstract

Management of agricultural pests requires an understanding of pest species diversity, their interactions with beneficial insects and spatial-temporal patterns of pest abundance. Invasive and agriculturally important insect pests can build up very high populations, especially in cropping landscapes. Traditionally, sampling effort for species identification involves small sample sizes and is labour intensive. Here, we describe a multi-primer high throughput sequencing (HTS) metabarcoding method and associated analytical workflow for a rapid, intensive, high-volume survey of pest species compositions. We demonstrate our method using the taxonomically challenging *Bemisia* pest cryptic species complex as examples. The whiteflies *Bemisia* including the *‘tabaci’* species are agriculturally important capable of vectoring diverse plant viruses that cause diseases and crop losses. Our multi-primer metabarcoding HTS amplicon approach simultaneously process high volumes of whitefly individuals, with efficiency to detect rare (i.e., 1%) test-species, while our improved whitefly primers for metabarcoding also detected beneficial hymenopteran parasitoid species from whitefly nymphs. Field-testing our redesigned *Bemisia* metabarcoding primer sets across the Tanzania, Uganda and Malawi cassava cultivation landscapes, we identified the sub-Saharan Africa 1 *Bemisia* putative species as the dominant pest species, with other cryptic *Bemisia* species being detected at various abundances. We also provide evidence that *Bemisia* species compositions can be affected by host crops and sampling techniques that target either nymphs or adults. Our multi-primer HTS metabarcoding method incorporated two over-lapping amplicons of 472bp and 518bp that spanned the entire 657bp 3’ barcoding region for *Bemisia*, and is particularly suitable to molecular diagnostic surveys of this highly cryptic insect pest species complex that also typically exhibited high population densities in heavy crop infestation episodes. Our approach can be adopted to understand species biodiversity across landscapes, with broad implications for improving trans-boundary biosecurity preparedness, thus contributing to molecular ecological knowledge and the development of control strategies for high-density, cryptic, pest-species complexes.

## Introduction

The ecology of a species is linked by definition to its species status. It underpins our understanding of species diversity in conservation genetics, biosecurity preparedness and developing effective pest management strategies. The correct identification of cryptic species can be challenging and often involves the use of molecular approaches. Timely identification and characterisation of pest species will greatly assist the development of effective management strategies to minimise impacts of crop losses due to pests on food security. Furthermore, techniques that enable the rapid screening and accurate identification of large numbers of individuals can accelerate research on a target pest species and lead to important insights into species ecology and biology.

The use of the maternally-inherited mitochondrial cytochrome c oxidase subunit I (mtCOI) partial gene 5’ (N) terminal region (i.e., typically one of 648 bp, 650 bp, or 658 bp) in ‘DNA barcoding’ via the Sanger sequencing method has contributed to molecular diagnostics of species (e.g., Hebert, Cywinska, Ball, & DeWaard, 2003; Ward, Zemlak, Innes, Last, & Hebert, 2005; Berry et al., 2004; Boykin et al., 2007; Dinsdale, Cook, Riginos, Buckley, & De Barro, 2010), and provided insights into the potential species diversity where cryptic species may co-exist (e.g., Hebert et al., 2003; Rao, Liew, Yow, & Ratnayeke, 2018; Tay, Beckett, & De Barro, 2016). Cryptic species such as the hemipteran whitefly *Bemisia tabaci* and related *‘non-tabaci’* species are excellent examples where putative species identification can be achieved via molecular characterisation of the partial mtCOI gene (e.g., Lee, Park, Lee, Lee, & Akimoto, 2013; Mugerwa et al., 2018), provided that PCR artefacts, such as NUMTs and/or poor quality sequence trace files and/or gene regions, were first removed (e.g., Elfekih et al., 2019; Kunz, Tay, Elfekih, Gordon, & De Barro, 2019; Tay, Elfekih, Court, et al., 2017; Vyskočilová, Tay, van Brunschot, Seal, & Colvin, 2018). Within the *B. tabaci* cryptic species complex, over 34 putative *B. tabaci* cryptic species have been reported (Kunz, Tay, Elfekih, et al., 2019; Lee et al., 2013; Mugerwa et al., 2018; Vyskočilová et al., 2018). The cryptic species status of the *B. tabaci* complex reflected the significant challenges associated with the correct identification of species status (reviewed by Boykin et al., 2018; De Barro, Liu, Boykin, & Dinsdale, 2011; Tay & Gordon, 2019). Species of the *B. tabaci* complex are notorious around the world as effective vectors of viruses that cause significant diseases of crop plants (Polston, De Barro, & Boykin, 2014). In east Africa, *B. tabaci* species transmit diverse viruses that cause diseases in cassava. Cassava mosaic disease (CMD) and cassava brown streak disease (CBSD) have led to significant crop losses and loss of production by smallholder farmers (Legg et al., 2011; Legg, Shirima, et al., 2014). Understanding the *Bemisia* species present in east Africa, and their patterns of abundance and distribution in different farming landscapes, is a necessary first step to managing this pest problem (Macfadyen et al., 2018).

In African nations such as Tanzania, Malawi and Uganda, cassava is typically grown by smallholder farming families, surrounded by non-cassava crops such as tomatoes, sweet potatoes, groundnuts, soybean, and maize. Native ecosystems that include unmanaged grazing lands, weeds, and shrubs, are also present and contain plants which are capable of supporting a diversity of whitefly species. Some may be novel species, which are yet to be characterised by morphological, behavioural and molecular approaches (e.g., Mugerwa et al., 2018; Vyskocilova, Seal, & Colvin, 2019; Vyskočilová et al., 2018). Globally, cassava is an important carbohydrate source for over 500 million people (<u>ARC</u> 2014, accessed 21-April-2019), and is planted in Africa, South America, and Asia/South East Asia. Thirteen of the top 20 global cassava producers are African countries (i.e., Nigeria, Angola, Ghana, Democratic Republic of Congo, Malawi, United Republic of Tanzania, Cameroon, Mozambique, Benin, Sierra Leone, Madagascar, Uganda and Rwanda (<www.worldatlas.com> up-dated 25-April-2017; accessed 21-April-2019). In other countries such as Thailand, Vietnam and Cambodia, it is grown as a source of high-quality starch and is used in diverse food products (Delaquis et al., 2018; Gotz & Winter, 2016; Graziosi et al., 2016; Kumar, Chang, Narayanan, & Ramasamy, 2017). Therefore, having techniques that enable rapid and cost-effective surveillance to investigate pest species compositions especially in cassava (and other food crops, i.e., tomatoes; other cash crops, i.e., cotton, ornamental plants) production landscapes are incredibly valuable. While identifying cryptic species diversity can best be achieved through molecular characterisation of appropriate genomic regions, adapting the concept of high-throughput sequencing (HTS) platforms to answer questions around species diversity and evolutionary interactions can be challenging.

The HTS platform (e.g., Illumina’s iSeq100, MiniSeq, MiSeq, NextSeq 550, NextSeq 1000 & 2000, NovaSeq 6000; Oxford Nanopore Technologies (ONT) range i.e., Flongle, MinION, GridION, and PromethION; PACBIO’s Single Molecule, Real-Time (SMRT) Sequencing and Sequel System/Sequel II System/Sequel IIe System) offers a range of scalable solutions (i.e., to best suit the research institute’s and target country’s infrastructure) for surveillance of insect species (e.g., see Hebert et al. 2018; Srivanthsan et al. 2021), including cryptic species that reach high population densities, provided that methods are sufficiently flexible to accommodate study designs tailored to address targeted research questions. For example, understanding host-parasitoid relationships, cryptic species diversity, host-plant relationships, and identifying keystone species are fundamental ecological questions common to diverse ecosystems. Addressing these ecological and evolutionary questions can be challenging and requires scaling up of the data collection processes. HTS platforms represent a solution when combined with appropriate species diversity databases, flexible bioinformatics and associated analytical pipelines, and optimised molecular methods.

The metabarcoding approach that incorporates the power of HTS platform and the utility of the DNA barcoding gene region aims to provide species identification at both population and community levels (i.e., not aimed at characterising all known mtCOI haplotypes at the individual level). In the cryptic *Bemisia tabaci* species complex however, molecular diagnostics is particularly susceptible to sub-optimal primer efficacies (Elfekih, Tay, et al., 2018; Mugerwa et al., 2018; Shatters, Powell, Boykin, Liansheng, & McKenzie, 2009; Tay, Elfekih, Court, et al., 2017). Partial or inconsistent Polymerase Chain Reaction (PCR) amplification, especially when targeting co-extracted DNA of individual cryptic species that may be found together, can lead to inaccurate conclusion of species composition within the surveyed agroecosystems. For example, a minor species may become over-represented within a population if co-amplified within a pool of major species but enjoys superior primer efficacies compared with the major species. Also, given the time, effort and cost necessarily associated with the preparation of HTS DNA libraries, especially where accurate input genomic DNA concentration is needed, metabarcoding should incorporate a ‘back-up’ step. Such a step may involve a second set of primers, ensuring the amplification of the same population, should the first set of primers fail and thereby avoiding a total PCR failure. Such a back-up design also enables capturing of species that are missed by the first primer pairs, and vice versa, or where longer DNA region is needed to improve molecular diagnostics of cryptic species complex. Provided that the primer pairs developed were sufficiently robust, within a pest insect species complex such as in the whitefly cryptic species, beneficial insects such as parasitoid species can also be investigated with appropriate sampling of pest-insect life stages (e.g., whitefly nymphs).

Here, we describe the HTS metabarcoding method and primers developed for surveying the *Bemisia* cassava whitefly, related *B. tabaci* cryptic species and associated parasitoid species complex, and the workflow for processing the large volume of HTS amplicon sequence reads generated. Our goal was to develop a multi-primer metabarcoding technique that enabled us to determine the dominant *Bemisia* species in any cassava field quantitatively, using a robust sample size in terms of numbers of individuals. We provide a verification of primer efficacies and both successful and failed species delimitation rates of the multi-primer method, and confirmed the method and primer efficacies via Sanger sequencing of individual field-sampled novel whiteflies species. Our samples were collected from geographically diverse cassava cropping landscapes in Uganda, Malawi and Tanzania, so test the limitations and usefulness of this approach especially for countries with significant infrastructure and economic challenges for researching agricultural pests impacting food security.

## Material & Methods

### Samples used in PCR primer efficacy tests

The DNA of whitefly cryptic species from Africa, Asia, South America, and Australia were individually extracted using the Qiagen Blood and Tissue DNA extraction kit (Cat #. 69506). Extracted DNA of all samples to be used for testing of PCR primer efficacies were individually eluted in 27.5 μL of EB. Whitefly species status was confirmed by PCR and Sanger sequencing using the primer set wfly-COI-F1/R1 and/or wfly-COI-F2/R2 (Suppl. Material I). Primer development and Sanger sequencing followed the methods of Elfekih, Tay, et al. (2018). Sanger sequencing was carried out at the BRF at JCSMR ANU, Canberra, ACT. Whitefly species used to verify primer efficacies and their respective GenBank accession numbers are listed in Table 1.

**Table 1:**
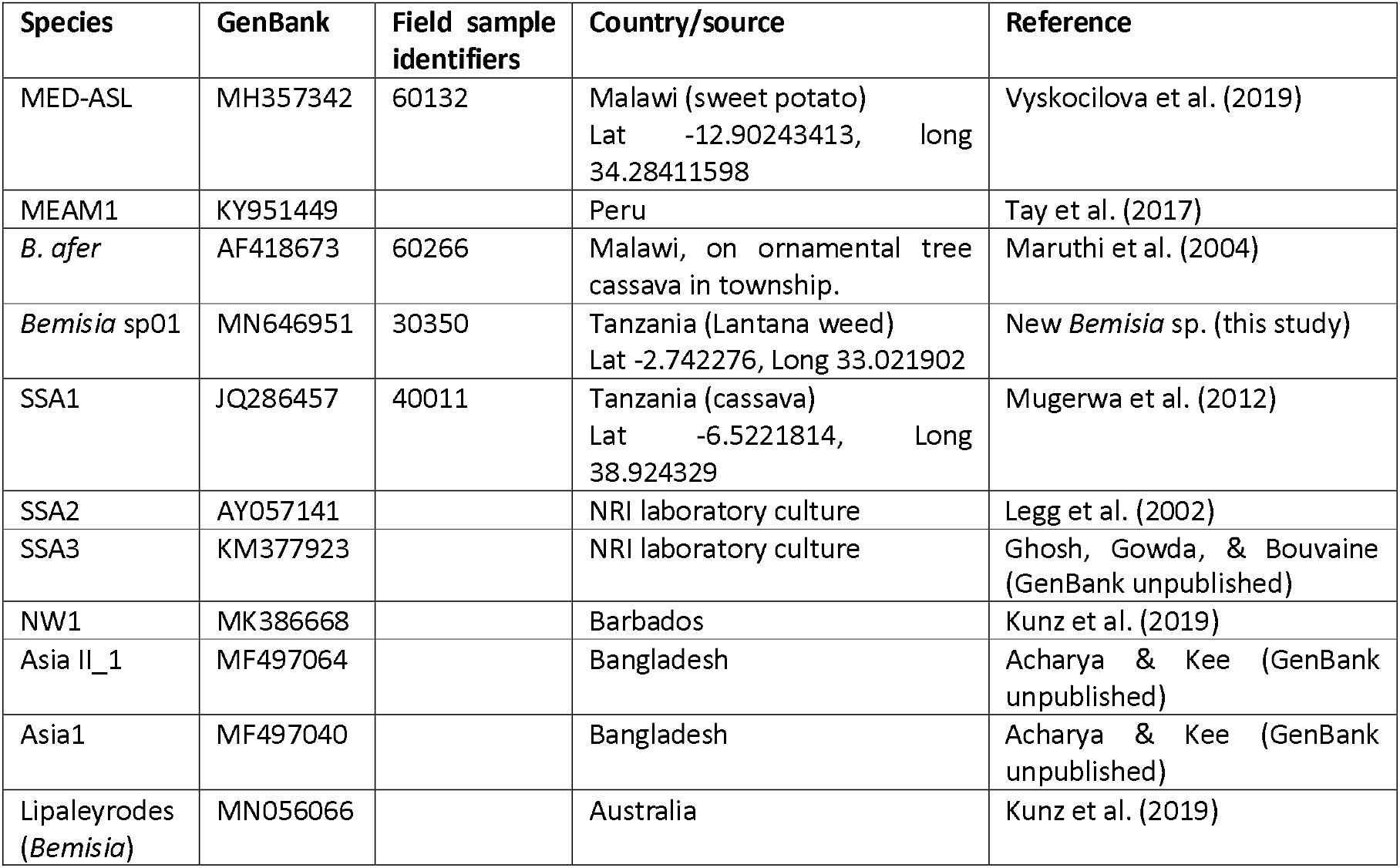
*Bemisia* species used to confirm PCR primer efficacies. With the exception of MN646951, all GenBank accession numbers provided represent partial mtCOI sequences of *Bemisia* species tested where 100% sequence identity against publicly available sequences have been detected. **Note:** KY951449 and MK386668: nucleotide position: 782-1438.

To ascertain PCR efficacies of the redesigned metabarcoding primers, various hypothetical field scenarios were simulated. These scenarios involved pooling of different genomic DNA (gDNA) that belonged to 11 separately extracted individuals of *Bemisia* species (Table 2) with predetermined species status based on the standard partial mtCOI dataset (Kunz, Tay, Elfekih, et al., 2019). These hypothetical scenarios included: (I) a two-species co-occurrence scenario, in which there is a dominant *Bemisia* whitefly species (e.g., SSA1) and a minor species present in different predetermined proportions (i.e., Table 2: Mixed-Ia (5%), Mixed-Ib (2.5%), and Mixed-Ic (1%)), and (II-IV) scenarios assuming co-occurrence of multiple mixed *Bemisia* cryptic pest species at different proportions based on estimates of gDNA concentration. With the exception of the test scenario ‘Mixed-III’ and ‘Mix-IV’ which were each replicated twice, all remaining test scenarios were replicated three times to estimate error rates and ensure protocol robustness.

rtain primer efficacies and species detection sensitivity by high-throughput sequencing platform. Average ± standard deviation of detection sensitivity (i.e., detection of observed proportion of taxa) for each category is provided based on amplicon reads as generated. Others’ include amplicons associated with NUMTs (e.g., MEAM2; Tay, Elfekih, Court, et al., 2017), where the MEAM1 species was used in the Mixed-Ic and Mixed-IV scenarios), unknown *Bemisia* genomic regions, parasitoids (*Encarsia*, *Eretmocerus*) partial mtCOI gene region, bacteria including secondary symbionts, fungal, and of unknown origins. ‘O’: Species not considered in test scenarios.

| Detection sensitivity | Selected known *Bemisia* species |  |  |  |  |  |  |  |  |  |  | Others† |
| --- | --- | --- | --- | --- | --- | --- | --- | --- | --- | --- | --- | --- |
|  | SSA1 | SSA2 | SSA3 | *Bemisia*. Sp01 (MN646951) | MEAM1 | MED | NW1 | Asia1 | AsiaII-1 | *Bemisia* sp. (MN056066) | *B. afer* |  |
| Mixed-Ia (1:20) | 95<br>(88.9±0.3) | 0 | 0 | 0 | 0 | 5<br>(6.1±0.7) | 0 | 0 | 0 | 0 | 0 | 0<br>(5±0.01) |
| Mixed-Ib (1:40) | 97.5<br>(94.8±0.3) | 0 | 0 | 0 | 0 | 2.5<br>(1.6±0.1) | 0 | 0 | 0 | 0 | 0 | 0<br>(3.6±0.2) |
| Mixed-Ic (1:100) | 99<br>(92±0.7) | 0 | 0 | 0 | 1<br>(1.2±0.4) | 0 | 0 | 0 | 0 | 0 | 0 | 0<br>(7.5±1.3) |
| Mixed-II | 0 | 0 | 0 | 0 | 0 | 0 | 25<br>(11.4±0.7) | 25<br>(12.7±0.9) | 25<br>(28±1) | 25<br>(20.2±1.2) | 0 | 0<br>(27.9±2.1) |
| Mixed-III | 0 | 0 | 0 | 0 | 0 | 0 | 30<br>(24.1±0.6) | 50<br>(30.3±0.4) | 10<br>(8.8±0.5) | 10<br>(14±0.4) | 0 | 0<br>(24.1±1.4) |
| Mixed-IV | 58<br>(42.5±4.9) | 8.8<br>(12.5±1.8) | 8.8<br>(10.5±0.7) | 6<br>(3±1.8) | 6<br>(7±0.6) | 6<br>(3.1±0.7) | 0 | 0 | 0 | 0 | 6<br>(2.9±0.6) | 0<br>(18.2±3.1) |

**Note:** † Only the first 10-15 amplicon contigs were searched for sequence identity by nucleotide blast (blastn) (i.e., optimised for somewhat similar sequences) search against the non-redundant (nr) NCBI DNA database. Verification of sequence identity was accomplished using the sanitized nucleotide database of Kunz, Tay, Elfekih, et al. (2019) as subject sequences, and using the NCBI’s Basic Local Alignment Search Tool (BLAST) “blastn suite-2sequences” option and using only top hits (i.e., those with 97-100% sequence identity). PCR primer amplification efficacies were ascertained on species representing significantly divergent evolutionary lineages (Kunz, Tay, Elfekih, et al., 2019) that included a novel *Bemisia* species (MN646951; From Tanzania field code: 30350, see Macfadyen et al., 2021), *Lipaleyrodes* (*Bemisia*) from Queensland, Australia (MN056066), *B. afer*, the African cassava whitefly species (SSA1, SSA2, SSA3), and species within the *B. tabaci* cryptic complex from diverse geographic origins (e.g., New World 1 (NW1); Mediterranean (MED), Middle East Asia-Minor 1 (MEAM1); Asia II-1, and Asia 1).

### Field samples survey methods

Cassava fields in Uganda, Tanzania, and Malawi were surveyed for *Bemisia* cryptic species between 2015-2016 (Fig. 1). In total, we conducted three data collection trips across a two-year period (trip 1, 1/8/2015 - 26/08/2015, trip 2, 5/4/2016 - 25/4/2016, trip 3, 29/10/2016 - 9/11/2016). In each region, we selected up to 10 cassava fields to survey and sample for nymph and adult *Bemisia* whiteflies (for sampling details see also Macfadyen et al., 2021). Briefly, up to *ca*. 50 adults were collected using an aspirator from the top leaves of cassava plants of known variety in four Malawi field sites (Table 2). Nymphs were collected from all sites by selecting leaves that were lower down on the plant (and so had visible nymphs in the second to forth instar age range on the underside of the leaf). The top lip of two centimetre diameter circular plastic vial (5ml volume) was used to cut small leaf discs that contained nymphs. These discs were placed in the vial filled with 99% ethanol. As a comparison, nymphs from other crops surrounding the focal cassava field were also collected. All ethanol-preserved samples were stored in −20°C prior to being processed in the laboratory, where nymphs were dislodged from leaves and placed in fresh Eppendorf (1.5mL) test tubes in batches of between 20 and 40 individuals per field (note that in some cases, fields sampled for *Bemisia* contained <20 individuals). For our research question we selected nymphs that had no obvious signs of parasitism by hymenopteran parasitoids in order to optimize the *B. tabaci* DNA in the sample. We randomly selected 26 field-collected *Bemisia* populations from Malawi, Uganda and Tanzania for analysis to demonstrate the efficacies of the HTS mtCOI amplicon approach under field conditions (Table 3).

**Fig. 1:**
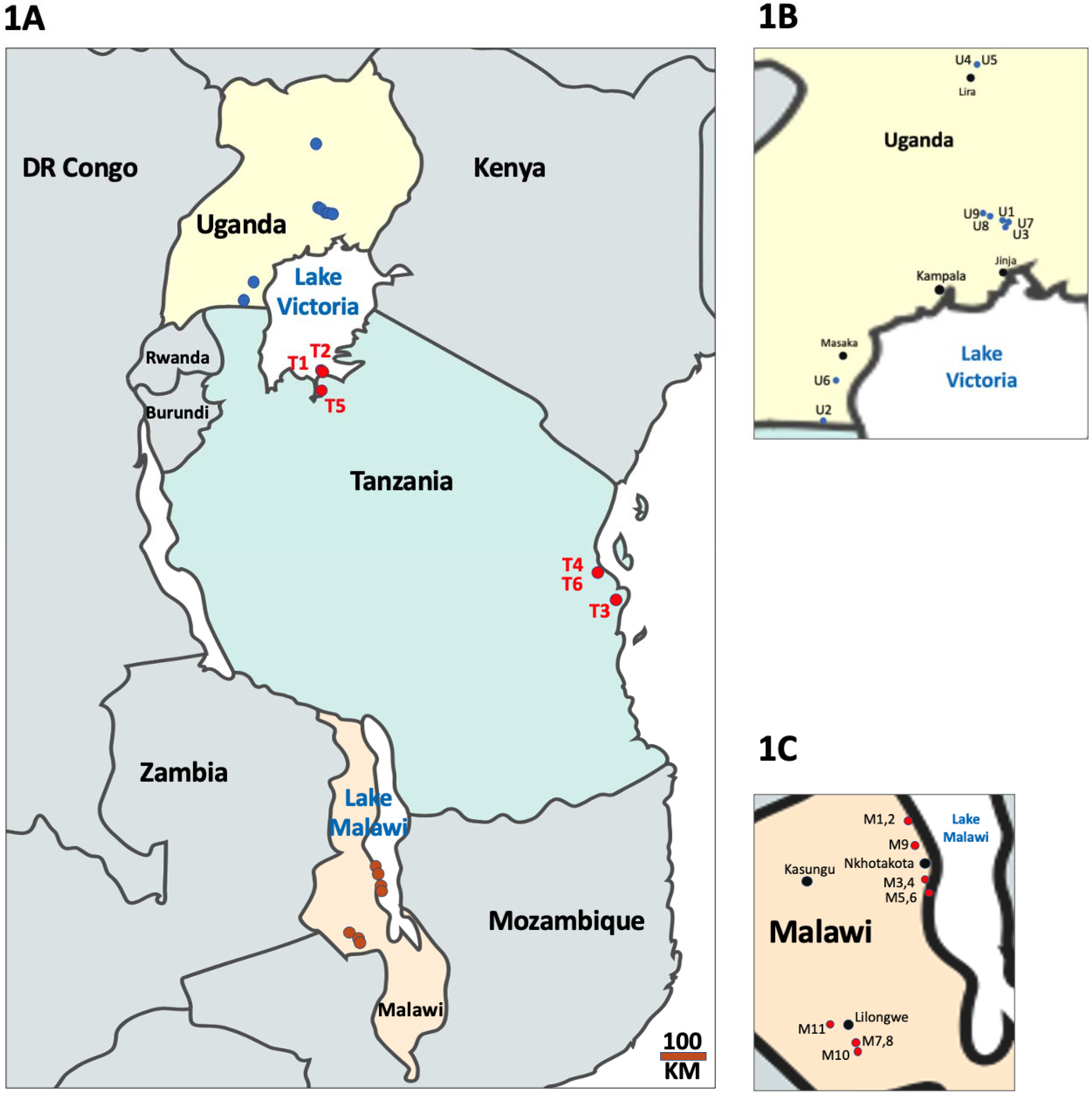
(A) *Bemisia* cryptic species sampling sites from Uganda, Tanzania and Malawi, with field site codes from Tanzania (i.e., T1 – T6) provided in (1A), Uganda (i.e., U1 – U9) in panel 1B, and Malawi (i.e., M1 – M11) in panel 1C. GPS co-ordinates of each sampling site are provided in Table 3. Maps of Africa produced using Mapchart <https://mapchart.net>.

**Table 3:** Twenty-six field-collected whitefly samples from Malawi, Uganda and Tanzania analysed by high-throughput amplicon sequencing. Life stages were nymphs (N); and adults (A). Sample size (n). Detected individuals for each species were estimated from proportions of amplicons. *Bemisia* cryptic species detected were: *B. tabaci* Indian Ocean (IO), Mediterranean (MED), *B*. sp. unknown (U), *B. afer* (Baf), *B*. sp. ‘Uganda’ (Ug1), African cassava whitefly sub-Saharan Africa 1 (SSA1), sub-Saharan Africa 2 (SSA2), *Trialeurodes* sp. (T), *Encarsia* spp. (Enc), *Eretmocerus* spp. (Eret), Bacterial (Bac). Nuclear mitochondrial sequences/PCR artefacts (NUMTs) were detected in all cases. GenBank accession numbers of all novel *Erectmocerus* and *Encarsia* species were MN646915-MN646927 and MN646928-MN646950 respectively, GenBank accession numbers of all novel *Bemisia* and *Trialeurodes* whitefly species detected were MN660053-MN660056 and MN660057-MN660058, respectively. Raw HTS amplicon reads can be downloaded from GenBank (SRA xxxx01, xxxx02, xxxx03). Site codes are as shown in Fig. 1.

| Country | Site ID<br>(site code) | Life stage | Host Plants (variety) | lat | long | Species composition | Total<br>reads<sup>#</sup> | Reads Identified<br>(%) |
| --- | --- | --- | --- | --- | --- | --- | --- | --- |
| Malawi | 60109/60121<br>(M1) | N, n= 26 | Cassava (Gomani) | -12.639611255 | 34.17544665 | SSA1: 12.7, Baf: 12.8, Enc: Y,<br>NUMTs: 0.4 | 43,652 | 43,383 (99.35%) |
|  | 60109/60121<br>(M2) | A, n= 7 | Cassava (Gomani) | -12.639611255 | 34.17544665 | SSA1: 6.4, Baf: 0.4, NUMTs: 0.01 | 54,392 | 54,135 (99.53%) |
|  | 60013/60126<br>(M3) | N, n=22 | Cassava (Gomani) | -13.03129506 | 34.29746802 | SSA1: 21.7, NUMTs: 0.17 | 68,562 | 68,224 (99.51%) |
|  | 60013/60126<br>(M4) | A, n= 8 | Cassava (Gomani) | -13.03129506 | 34.29746802 | Baf: 7.84, NUMTs: 0.09 | 44,424 | 44,166 (99.42%) |
|  | 60127/60250<br>(M5) | N, n= 20 | Cassava (Gomani) | -13.11325369 | 34.32143017 | SSA1: 5.1, Baf: 14.5, NUMTs: 0.07 | 34,660 | 32,169 (92.81%) |
|  | 60127/60250<br>(M6) | A, n= 20 | Cassava (Gomani) | -13.11325369 | 34.32143017 | SSA1: 15.9, Baf: 3.8, NUMTs: 0.13,<br>Bac: 0.01 | 62,600 | 62,193 (99.35%) |
|  | 60032/60263<br>(M7) | N, n= 18 | Cassava (Manyokola) | -14.11054382 | 33.81826233 | SSA1: 0.63, Baf: 14.7, U: 0.08,<br>NUMTs: 1.45 | 43,156 | 40,341 (93.48%) |
|  | 60032/60263<br>(M8) | A, n= 15 | Cassava (Manyokola) | -14.11054382 | 33.81826233 | SSA1: 5.3, Baf: 5.9, U: 0.56, NUMTs:<br>2.6 | 72,172 | 66,784 (92.53%) |
|  | 60010/60242<br>(M9) | N, n= 30 | Cassava (Beatrice) | -12.804086285 | 34.21788623 | SSA1: 18.7, Baf: 10.1, NUMTs:<br>0.91, Enc: Y, Eret: Y | 63,506 | 62,888 (99.03%) |
|  | 60037/60265<br>(M10) | N, n= 40 | Cassava (Manyokola) | -14.1593299559021 | 33.8348985175844 | SSA1: 0.6, Baf: 38.7, Enc: Y, NUMTs:<br>0.16 | 44,808 | 44,228 (98.71%) |
|  | 60039/60040<br>(M11) | N, n= 40 | Cassava (Manyokola) | -13.9888087453993 | 33.6391004272323 | SSA1: 38.4, Baf: 0.7, Eret: Y, T: 0.2,<br>NUMTs: 0.52 | 36,530 | 36,207 (99.12%) |
| Uganda | 10109/10230<br>(U1) | N, n= 40 | Cassava (TME_14) | 0.93830798 | 33.16688609 | SSA1: 32.8, SSA2: 3.4, Eret+Enc: Y<br>(0.5), NUMTs: 2 | 49, 601 | 47,906 (96.58%) |
|  | 20018/20019 (U2) | N, n= 40 | Cassava (Omongole) | -0.98750407 | 31.42085652 | **SSA1:** 32.8, **Baf:** 2.5, **Eret+Enc:** Y (4.1), **NUMTs:** 0.3 | 38,082 | 37,752 (99.13%) |
|  | 10085/10336 (U3) | N, n= 40 | Cassava (TME_14) | 0.87832027 | 33.19462426 | **SSA1:** 35.6, **Baf:** 0.13, **Eret+Enc:** Y (2.8), **NUMTs:** 0.96 | 68,176 | 67,185 (98.55%) |
|  | 43130 (U4) | N, n= 20 | Cassava (TME_14) | 2.35231436 | 32.93393845 | **SSA1:** 9.9, **SSA2:** 3.1, **Baf:** 2.4, **Eret:** Y, **NUMTs:** 1.8 | 7,486 | 6,531 (87.24%) |
|  | 43131 (U5) | N, n= 40 | Cow peaCowpea | 2.35246646 | 32.93482051 | **SSA1:** 8.1, **MED:** 21.8, **U:** 1.6, **Eret:** Y, **NUMTs:** 2.1 | 6,674 | 5,883 (88.15%) |
|  | 53371 (U6) | N, n= 40 | Sweet potato | -0.52519054 | 31.6441164 | **SSA1:** 1, **MED:** 26.2, **IO:** 0.4, **Ug1:** 11, **NUMTs:** 0.14, **Eret:** Y (0.1) | 58,646 | 56,426 (96.21%) |
|  | 10362 (U7) | N, n= 8 | Cassava (Nase_14) | 0.92017081 | 33.21595328 | **SSA1:** 5.1, **SSA2:** 1.6, **NUMTs:** 0.14 | 7,305 | 6,202 (84.90%) |
|  | 33026 (U8) | N, n= 20 | other crop, Malakwang Rosella-like | 0.97703406 | 33.04814937 | **SSA1:** 0.6, **MED:** 19.2, **Eret:** Y, **NUMTs:** 0.2 | 54,877 | 54,726 (99.72%) |
|  | 33042 (U9) | N, n= 20 | Cassava (Nase_14) | 1.00622674 | 32.98948722 | **SSA1:** 18.4, **MED:** 0.03, **Eret+Enc:** Y (1.3), **NUMTs:** 0.18 | 47,014 | 46,812 (99.57%) |
| Tanzania | 30145/30157 (T1) | N, n= 40 | Cassava (busanagulwa) | -2.388214545 | 33.05174421 | **SSA1:** 33, **Eret+Enc:** Y (6.5), **NUMTs:** 0.27 | 58,826 | 58,407 (99.29%) |
|  | 30052/30061 (T2) | N, n= 40 | Sweet potato in cassava field | -2.41603623925181 | 33.0826151071912 | **SSA1:** 39.3, **Baf:** 0.4, **Enc:** Y, **NUMTs:** 0.3 | 62,988 | 62,589 (99.37%) |
|  | 40013/40014 (T3) | N, n= 40 | Cassava (Kiroba) | -7.15118632229587 | 39.1807717492072 | **SSA1:** 31.5, **Baf:** 6, **NUMTs:** 1.1 | 71,134 | 68,629 (96.48%) |
|  | 40279/40288 (T4) | N, n= 40 | Cassava (Kiroba) | -6.58507491183717 | 38.8436025425142 | **SSA1:** 32.4, **Baf:** 1.1, **Eret+Enc:** Y (5.8), **NUMTs:** 0.34 | 49,610 | 49,154 (99.08%) |
|  | 30359 (T5) | N, n= 12 | Cassava (unknown) | -2.802858945 | 33.05936783 | **SSA1:** 9.5, **U:** 1.2, **NUMTs:** 0.1 | 6,931 | 6,219 (89.73%) |
|  | 70120 (T6) | N, n= 20 | Cassava (marfaransa) | -6.59705482 | 38.81119587 | **SSA1:** 14.2, **MED:** 0.1, **Eret+Enc:** Y, **NUMTs:** 4.3 | 6,029 | 5,912 (98.06%) |

**Note:** To calculate the estimated detected individuals for each species per sampling site, the proportion of amplicons assigned to each putative species was first determined by mapping to the reference species sequence (outlined in Fig. 2) and then divided by the total Illumina raw output fastq (i.e., Total reads<sup>#</sup>) reads from the pooled population.

### DNA extraction and HTS library preparation

DNA extraction of pooled individuals was carried out using the Qiagen Blood and Tissue DNA extraction kit (cat. #69056) and followed manufacturer’s instructions. Extraction of pooled individuals was life-stage specific (i.e., extraction of pooled adults or pooled nymphs were carried out separately). Extracted gDNA from each pooled sample was quantified for DNA concentration and standardised to 0.5 ng/μl prior to PCR amplification of target mtCOI region. Traditionally, molecular diagnostics for *Bemisia tabaci* cryptic species have relied on the F-C1-J-2195 and the TL2-N-3014 primers (Crozier, Crozier, & Mackinlay, 1989; Frohlich, Torres-Jerez, Bedford, Markham, & Brown, 1999; Roehrdanz, 1993; Simon et al., 1994) that amplified approximately 750bp of the 3’ COI terminal region. This primer set was developed based on Diptera, Lepidoptera, Coleoptera, and Hymenoptera but was not originally reported for Hemiptera ( Roehrdanz, 1993; Simon et al., 1994). Recent evaluations of these primers have found poor efficacies when applied in *B. tabaci* cryptic species (e.g., Elfekih, Tay, et al., 2018; Mugerwa et al., 2018) that contributed to the mis-identification of pseudo species (Kunz, Tay, Elfekih, et al., 2019; Tay, Elfekih, Court, et al., 2017). The *B. tabaci* cryptic species includes >34 well-defined species with interspecific nucleotide distances that ranged from *ca*. 3% to >18% (Dinsdale et al., 2010; Kunz, Tay, Elfekih, et al., 2019; Lee et al., 2013). To ensure maximal PCR efficacies of our primer sets, we aligned complete COI genes as obtained via HTS platform, from a range of species that included SSA1, SSA2 (Kunz, Tay, Elfekih, et al., 2019); MED cryptic species (Rossitto De Marchi, Kinene, Mbora Wainaina, Krause-Sakate, & Boykin, 2018; Tay, Elfekih, Court, et al., 2017; Vyskocilova et al., 2019); Asian species (Tay, Elfekih, Court, Gordon, & De Barro, 2016); *Bemisia* ‘JpL’ (Tay, Elfekih, Polaszek, et al., 2017), Australian (AUS), Indian Ocean, and Middle East Asian Minor 1 (MEAM1) species (Tay, Elfekih, Court, et al., 2017), to ascertain most conserved regions for designing and developing our two sets of complementary primers (wfly-PCR-F1/R1 and wfly-PCR-F2/R2; Suppl. Material 1). Primers were developed using Oligo 7 (Molecular Biology Insights, Colorado Springs, CO, USA) with minimal primer duplex and hairpin structure formation, with 55-65°C Tm, for expected amplicon size of 550-600bp, and with minimal false primer annealing sites.

Two PCR reactions were carried out for each pooled sample using primers wfly-PCR-F1/R1 and wfly-PCR-F2/R2 for the first and second reaction, respectively. For the PCR amplification of each pooled sample, we used 3.5ng gDNA template in a 35μL PCR reaction volume and 30 PCR cycles (PCR conditions described in Suppl. Material I). Amplicons from each of the pooled samples were ascertained for PCR amplification success on 1.25% 1× TAE agarose gel stained with GelRed and visualised on a UV-transilluminator. Amplicons were then purified using AMPure XP beads as instructed in the Illumina **16S Metagenomic Sequencing Library Preparation** (<u>Part # 15044223 Rev. B</u>) protocol and resuspended in 25 μL Qiagen EB (instead of the recommended 52.5μL). Purified pooled amplicons were individually quantified for DNA concentration using a Qubit v2.0 fluorometer (Life Technologies) and Qubit HS dsDNA assay kit and indexed by amplifying 5.0 ng of the purified amplicon in 50μL PCR volume over 11 PCR cycles (see Suppl. Material I). Indexed amplicons were purified using AMPure XP beads as described above and concentration ascertained using Qubit v2.0. Equal proportions of purified indexed amplicon (i.e., 100ng) from each pooled sample were mixed with 6x Loading Dye and run on 1.25% low melt agarose at 75 V for 2 hours. The F1/R1 combined amplicons and the F2/R2 combined amplicons were excised and purified using the Zymoclean Gel DNA Recovery Kit (cat. # D4001, D4002, D4007, D4008) following the manufacturer’s protocol. Purified indexed amplicons were quantified for double stranded DNA concentration using Qubit and fragment size was estimated using Tapestation (Agilent Technologies). Amplicons were calculated for their nano-molarity and diluted to 4 nM (see Illumina <u>470-2016-007-B</u>) for HTS run on MiSeq sequencer using the Illumina MiSeq Reagent Kit Version 3 (600bp pair-ended). A detailed protocol for the amplicon library preparation and related primer sequence information is provided as Suppl. Material I.

### HTS amplicon data processing

Fastq sequences representing individual populations post MiSeq runs were trimmed using Trimmomatic version 0.36 (Bolger, Lohse, & Usadel, 2014) by quality (Leading:3; Trailing: 3; Slidingwindow:4:15, minlength: 125bp) and imported into Geneious v11.1.5 as pair-ended reads of the two metabarcoding primer sets for simultaneous mapping to reference sequences. Amplicon sequence reads from each population were mapped to curated barcode reference sequences of representative *Bemisia* species (see below) using the Geneious Mapper option, and selecting to trim amplicon reads with options set to ‘Annotate new trimmed region, error probability limit set to 0.05, and trimmed off ≥23bp at both 5’ and 3’ ends. Customed mapping options within Geneious Mapper function were: no gaps allowed, word length set to 16; randomly map multiple best matches, index word length set at 15, maximum ambiguity option set to 8, and with two separate runs of 3% and 5% maximum mismatches per read for all *B. tabaci* species; 7% for *B. afer*, and 12% for parasitoids (Fig. 2). Unmapped reads after each round were used in mapping against other *Bemisia* and parasitoid reference sequences. A Geneious *de-novo* assembly step with default setting was carried out separately for each primer-pair, and only so when all reference whitefly and parasitoid sequences had been mapped and when un-mapped reads were typically low (i.e., ≤4,000 pair-ended reads). For the *de-novo* assembly, the top 10 most common contigs were determined for their sequence identity using nucleotide blast (blastn) within NCBI database. A schematic representation of the described workflow is provided in Fig. 2.

**Fig. 2:**
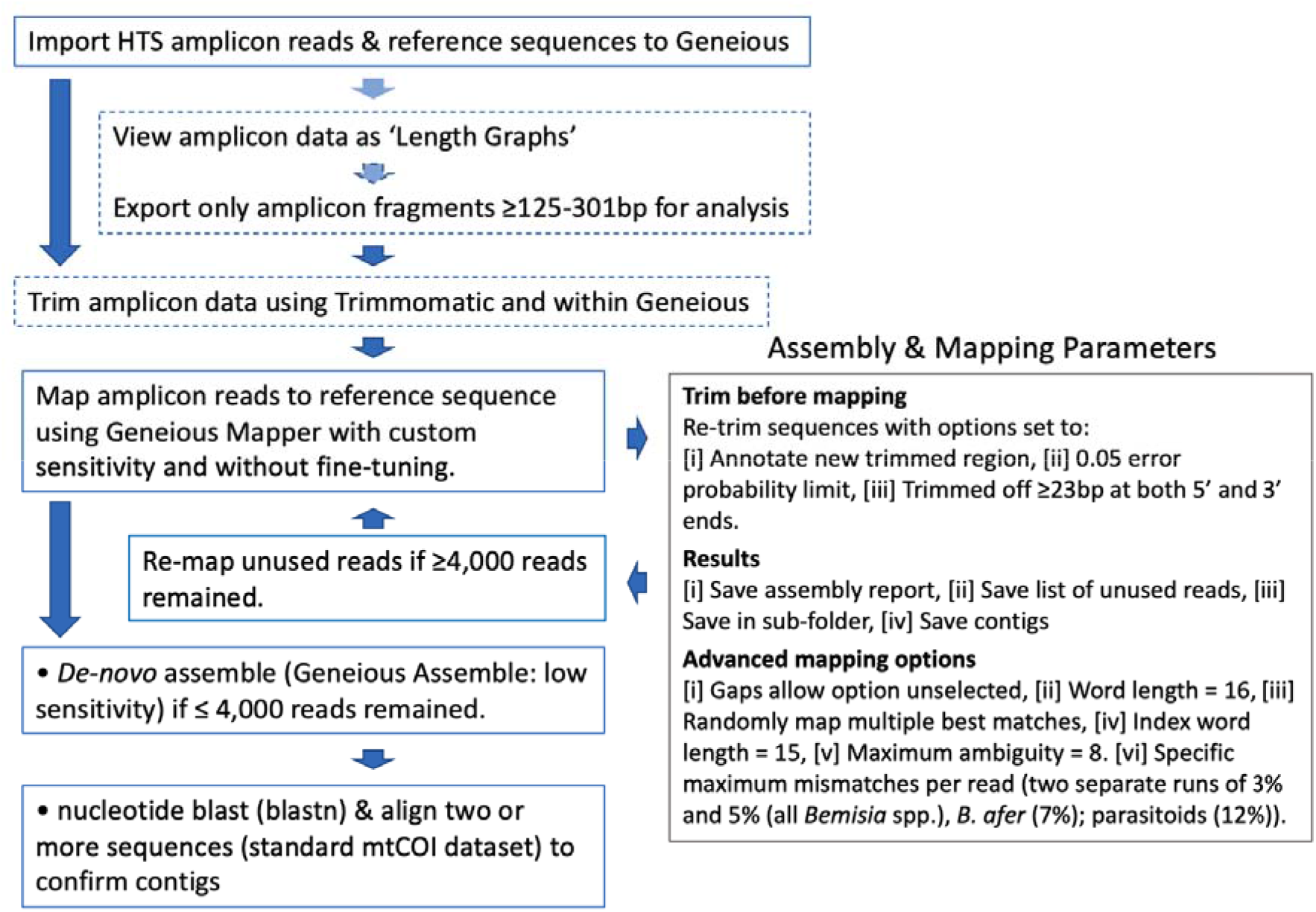
Workflow for analysis of MiSeq generated high-throughput sequencing of mtCOI amplicon reads for estimating proportions of *Bemisia* species in the African cassava cultivation landscape, including assessment of hymenopteran parasitoid species. Optional steps are indicated in dashed box depending on whether amplicon libraries were gel-purified (refer to Suppl. Material I) to remove small fragments. Quality of amplicons can also be improved if desired, by using an Illumina sequence trimming program such as Trimmomatic (Bolger et al., 2014). The *de-novo* assembly step enables novel *Trialeurodes*-, parasitoid- and bacterial-related sequences, as well as NUMTs, to be identified. NUMTs were identified through detection of INDELs, premature stop codons, and presence of unexpected amino acid residues at highly conserved COI regions following the methods of Kunz, Tay, Elfekih, et al. (2019).

### Reference sequences for identification of *Bemisia* mtCOI

Within the African cassava cultivation landscape context, the proportion of specific *Bemisia* and related cryptic species detected was estimated as represented by the proportion of total amplicon fragments. Our reference mtCOI sequences were from the following *Bemisia* species: SSA1 (GenBank JQ286457, nt 59-715), SSA2 (GenBank AY057141, nt 29-685), SSA3 (GenBank KM377923, nt 20-676), MEAM1 (GenBank KY951449, nt 782-1438), IO (GenBank KY951448, nt 782-1438), *B. afer* (GenBank AF418673, nt 59-715), Uganda1 (GenBank KX570857, nt 34-690), ‘African’ MEAM2 (GenBank KX570778, nt 34-690), MED (GenBank KY951447, nt 782-1438), SSA6 (GenBank KX570852, nt 34-690), SSA9 (GenBank KX570856, nt 34-690), SSA10 (GenBank KX570804, nt 34-690), SSA13 (GenBank KX570833, nt 34-690) and novel sub-Saharan *Bemisia* species (i.e., MN646951; MN646952) detected by Sanger sequencing from field-surveyed populations (Macfadyen et al., 2021; this study). We confirmed the authenticity of these reference sequences as free from NUMTs following the methods of Kunz, Tay, Elfekih, et al. (2019).

### Parasitoids reference sequences

*Eretmocerus* and *Encarsia* parasitoid mtCOI reference sequences from the sub-Saharan region surrounding our field survey sites were built up (MN646919, MN646949) during the course of the analysis workflow (see Fig. 2). Due to the anticipated significant whitefly hymenopteran parasitoid species diversity in the sub-Saharan region, we arbitrarily determine a cut-off value of 6% within the Geneious ‘Map to Reference’ function.

### Assessment of species delimitation using a stagged over-lapping amplicon approach

To assess efficacies of our two-primer set approach for metabarcoding of cryptic *Bemisia* and related whitefly species, we downloaded the clean *B. tabaci* mtCOI database of Kunz, Tay, Elfekih, et al. (2019) and performed phylogenetic analyses based on 315bp aligned nucleotides at 50bp sliding window sizes for primer pairs ‘wfly-PCR-F1/R1 (i.e., 3’ mtCOI barcode nucleotide position (nt) 1-315; nt 51-365; nt 101-415; nt 151-465), and ‘wfly-PCR-F2/R2’ (i.e., nt 140-454; nt 190-504; nt 240-554; nt 290-604; nt 340-654), as well as for the full length of wfly-PCR-F1/R1 amplicon (473bp) and the wfly-PCR-F2/R2 amplicon (518bp), and compared that with the species delimitating powers based on uncorrected pair-wise nucleotide similarity (i.e., p-dist) distance method of the full 657bp 3’ mtCOI *B. tabaci* barcoding gene. Sequence alignment using MAFFT Alignment (selecting ‘Auto’ option for the Algorithm setting, 200PAM / K=2 scoring matrix, and gap open penalty and offset value set at 1.53 and 0.123, respectively) (Katoh & Standley, 2013) and trimming were carried out in Geneious v11.1.5. Confidence of clustering of identified species was estimated using Ultrafast bootstrap as implemented in IQ-Tree, and followed the methods as described in Kunz, Tay, Elfekih, et al. (2019). Briefly, aligned and trimmed sequences were imported as fasta formatted file and imported to the IQ-Tree website <http://iqtree.cibiv.univie.ac.at> (Trifinopoulos, Nguyen, von Haeseler, & Minh, 2016). The ‘Automatic’ substitution model selection option was selected and we specified 1,000 ultrafast bootstrap approximation (Minh, Nguyen, & von Haeseler, 2013) with 1,000 maximum iterations and 0.99 as the minimum correlation coefficient. Visualisation and assessment of correct clustering of species were carried out using FigTree v1.4.4 (2006-2018 Andrew Rambaut, Institute of Evolutionary Biology, University of Edinburgh).

## Results

### Primer efficacies

The primers wfly-COI-F1/R1 and/or wfly-COI-F2/R2 were shown to be highly efficient in successful PCR amplification of diverse representative *Bemisia* species that included Australian species (Lipaleyrodes (*Bemisia*)), Asian, New World (NW), Mediterranean (MED), Middle East Asian Minor (MEAM1), sub-Saharan African (SSA) species, and *‘non-tabaci’* species (i.e., *B. afer*). Sanger sequencing of PCR amplicon from test *Bemisia* cryptic species (Table 1) partial mtCOI confirmed species identity when compared against both the updated standard *B. tabaci* mtCOI reference dataset and the uncorrected pairwise nucleotide distances (p-dist; see Kunz, Tay, Elfekih, et al., 2019). While Sanger sequencing confirmed sequence integrity such as through conservation of amino acid residues against published *B. tabaci* cryptic species HTS-derived full mtDNA genomes (e.g., Kunz, Tay, Court, et al., 2019; Tay, Elfekih, et al., 2016; Tay, Elfekih, Court, et al., 2017; Tay, Elfekih, Polaszek, et al., 2017; Vyskočilová et al., 2018), results from HTS amplicon analyses readily detected NUMTs such as those similar to ‘MEAM2’ (Delatte et al., 2005; Tay, Elfekih, Court, et al., 2017), as well as partial mtCOI genes with INDELs. Detection of co-amplified pseudogenes/NUMTs demonstrated the complexity of the *Bemisia* genome (Chen et al., 2016; Xie et al., 2017; Xie et al., 2018). HTS-amplicon analyses also shed light on the unexpected co-amplification of non-target whitefly genomic regions, as well as fungal, bacterial symbionts, and parasitoids genomic regions, and highlighted the complexity of trophic interactions in *Bemisia* whiteflies (e.g., Rao et al., 2018; Shamimuzzaman et al., 2019; Tay, Elfekih, Polaszek, et al., 2017).

### HTS-amplicon protocol and analysis workflow

Test scenarios for detection and identification of *Bemisia* cryptic species in predetermined proportions (i.e., 20:1; 40:1; 100:1) showed that the HTS large scale sampling protocol and the analysis workflow (Fig. 2) confidently identified the dominant as well as minor species in approximate expected proportions (Table 2). Similarly, in the hypothetical mixed-species composition scenario that included different proportions of mixed species (i.e., scenarios ‘Mixed-II, ‘Mixed-III, ‘Mixed-IV’; Table 2), the protocol also successfully detected all test species while the expected proportions were affected by low amount of co-amplified NUMTs/pseudogenes, bacterial, fungal, parasitoid, and non-specific whitefly genomic regions (Table 2). The test scenarios therefore provided significant insights into characteristics associated with our primers and the robustness of our multi-primer metabarcoding protocol. For example, across various scenarios, detected amplicon coverage regularly fell below the expected values due to non-specific PCR co-amplification of nuclear genome regions, endosymbionts, and environmental DNA contaminants.

### Field application of HTS-amplicon method

Applying the HTS multi-primer metabarcoding protocol to field-collected samples from Malawi, Tanzania, and Uganda showed that in the African cassava fields, differences in *Bemisia* species compositions existed (Kalyebi et al., 2018; Macfadyen et al., 2018; Macfadyen et al., 2021). A total of eight whitefly species (i.e., SSA1, SSA2, IO, MED, *B. afer*, Uganda 1, two novel *Bemisia* species), and a novel *Trialeurodes* species, were detected in different field sites (Table 3, Fig. 3). Multiple novel mtCOI haplotypes for *Eretmocerus* species (MN64915-MN64927) and *Encarsia* species (MN646929-MN646950) were also detected (Table 3). Parasitoid sequences were only ever detected in nymphal samples, and this was despite targeted exclusion of parasitised nymphal samples that could be visually identified prior to genomic DNA extraction. Detection of parasitoids is therefore likely the result of early stages of parasitism where visual identification would be most challenging. While our findings suggested high parasitism rates in the agricultural landscapes in Uganda, Tanzania, and Malawi, we nevertheless advise caution in interpreting the *Bemisia* parasitism rates, parasitoid species, and genetic diversity in these natural agricultural settings, as our primary experimental design was to develop a metabarcoding approach for surveys of the hemipteran *Bemisia* cryptic pest species only.

**Fig. 3:**
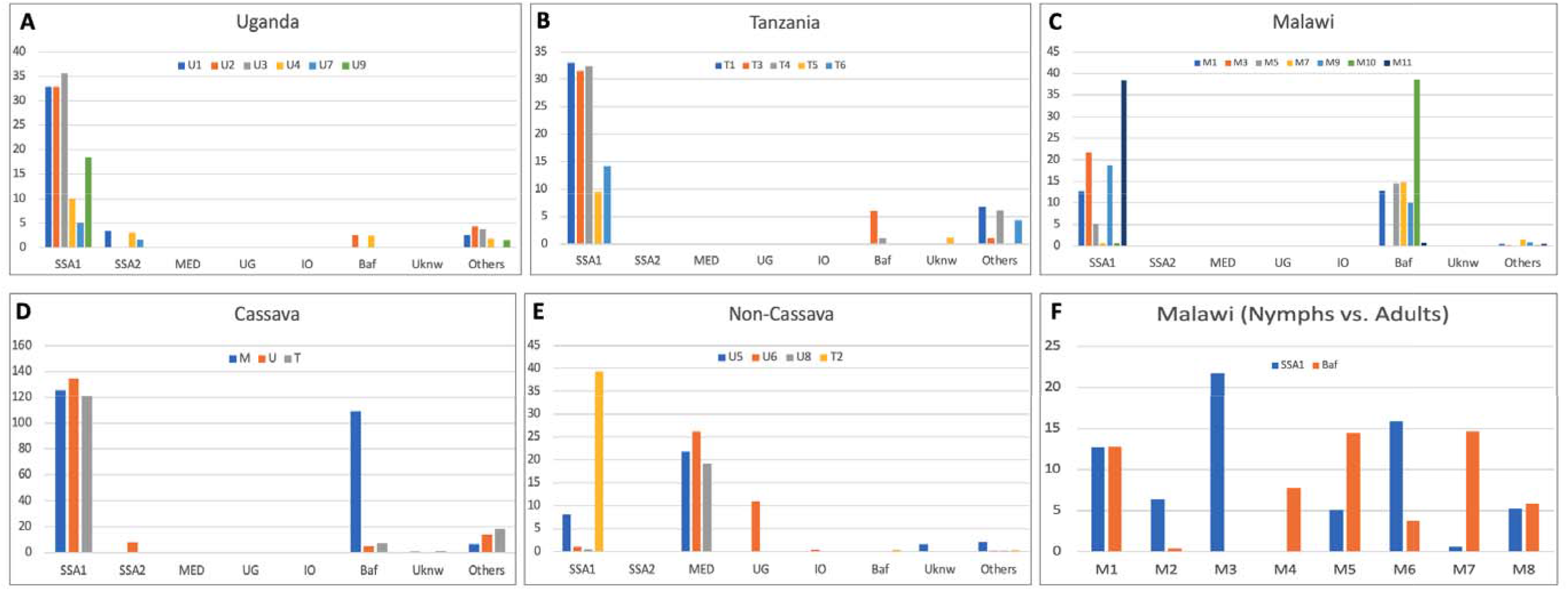
Summary results of *Bemisia tabaci* cryptic species from Uganda (panel A), Tanzania (panel B), and Malawi (panel C). Site codes are as provided in Table 3. *Bemisia* species are sub-Sharan Africa 1 (SSA1), sub-Saharan Africa 2 (SSA2), Mediterranean (MED), Uganda 1 (UG), Indian Ocean (IO), *B. afer* (Baf), and unknown (‘Uknw’) species from *Bemisia* and *Trialeurodes* genera. Low detection rates of COI-related pseudogene, parasitoid (*Eretmocerus* and *Encarsia* genera), and bacterial/fungal sequences are grouped in the ‘others’ category. Whitefly species compositions are also compared between cassava and non-cassava host plants (panels D and E, respectively), and between nymphal (M1, M3, M5, M7) and adult (M2, M4, M6, M8) life stages from four cassava sites in Malawi (panel F).

The sub-Saharan African 1 (SSA1) *Bemisia* species was identified as the dominant species in cassava fields from Uganda and Tanzania (Figs. 3A, 3B), while in Malawi both SSA1 and *B. afer* were present in approximately similar proportions (Fig. 3C). Overall, the SSA1 species was most prevalent in cassava landscape (Fig. 3D). We also detected the SSA1 species on non-cassava crop planted within a cassava field (‘T2’ in Fig. 3E; Table 3). The *B. tabaci* MED cryptic species complex was present as the dominant species in Uganda from three non-cassava host crop sampling sites (Fig. 3E; Table 3). Sweet potato crops also hosted other *Bemisia* species including Uganda 1 (‘UG’) and Indian Ocean (‘IO’), although both UG and IO species were not detected on sweet potato within cassava fields in Tanzania (‘T2’), and could suggest potential behavioural differences between SSA1 and UG/MED species (Fig. 3E) impacting on species-specific abundances. In Malawi, *B. afer* and SSA1 species dominated different field sites (Table 3, Figs. 3C, 3F). Their detection showed significant variability in species composition within the same field sites depending on the life-stage sampled (i.e., nymphal vs. adults; Fig. 3F). Field sample analyses also detected NUMTs and bacterial/fungal origins in the African populations, similar to our hypothetical scenarios (Table 2), thereby demonstrating consistencies between our HTS workflow and primer-efficacy assessments.

### Assessment of species delimitation efficacies between amplicon sizes

Short amplicons of between 130bp – 319bp (e.g., Brandon-Mong et al., 2015; Leray et al., 2013) but also longer amplicon lengths (e.g., 421bp, Hajibabaei, Porter, Wright, & Rudar, 2019) have been proposed for COI metabarcoding. Our assessment of species delimitation powers by short (i.e., 315bp) versus full amplicon lengths based on distance methods (i.e., p-dist) of our wfly-PCR-F1/R1 (473bp) and wfly-PCR-F2/R2 (518bp) primer sets, as well as their combined 657bp barcode gene length for the *B. tabaci* cryptic species complex showed that metabarcoding involving the full 657bp was needed to fully delimit between *Bemisia* species (Fig. 4). Species detection and delimitation involving the various shorter 315bp nucleotide lengths along the 657bp barcoding gene region often failed to differentiate certain cryptic species complexes such as those within the Mediterranean (MED), New World (NW), sub-Saharan Africa (SSA), and Australia (AUS) clades (Fig. 4). Interestingly, other invasive species such as the SSA2 (e.g., Hajibabaei et al., 2019) and the MEAM1 (De Barro et al., 2011; Elfekih, Etter, et al., 2018), as well as the Asia II-1 species that has recently emerged as important vector species in the transmission of the Sri Lankan Cassava Mosaic Virus (SLCMV) in South East Asia (Chi et al., 2020) could be readily identified by any of the 315bp COI region.

**Fig. 4:**
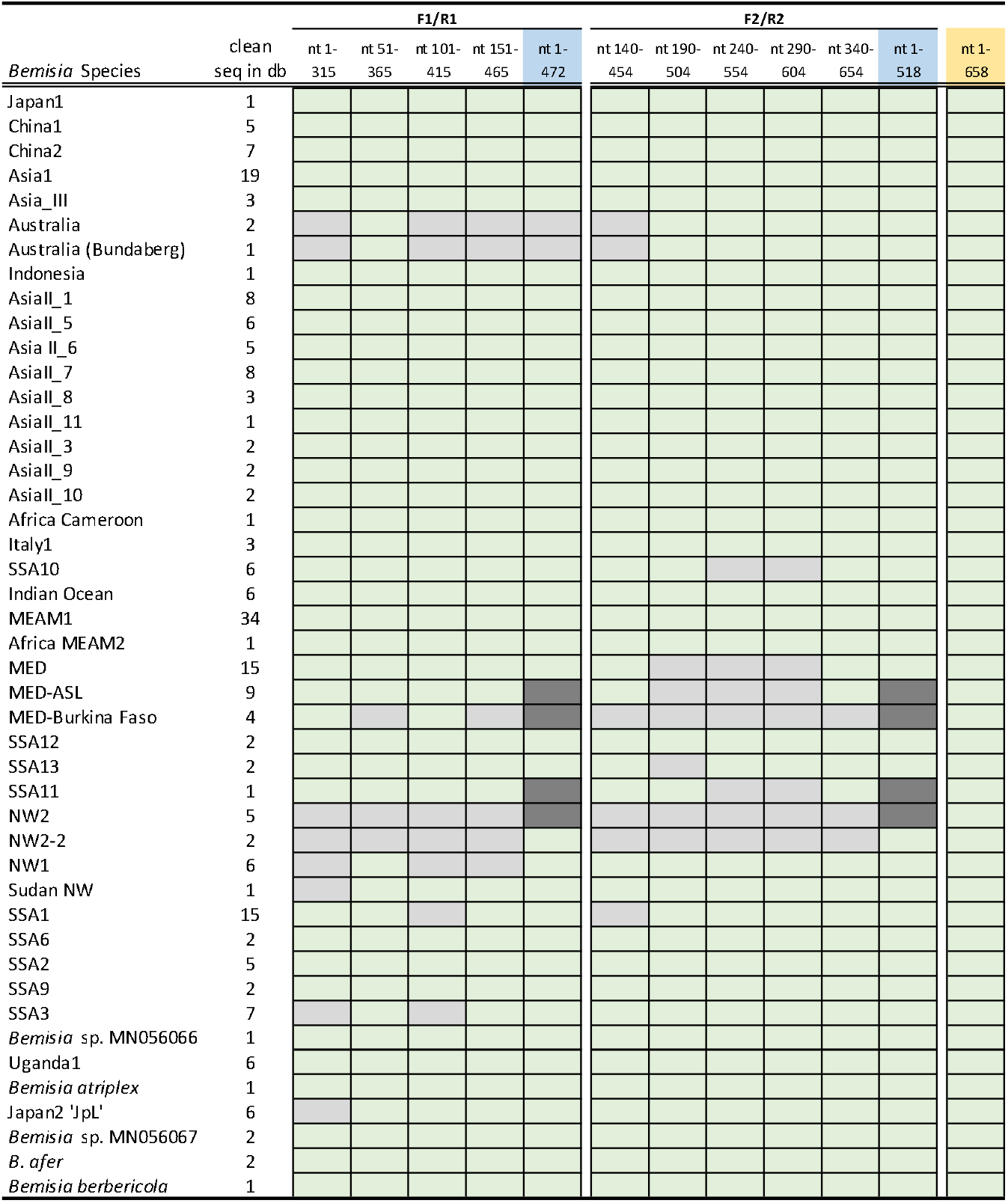
Summaries of efficacies of species delimitation powers based on the p-dist method across the 657bp 3’ mtCOI barcoding gene region in the cryptic *Bemisia tabaci* and *non-tabaci* species complex. The metabarcoding primer pair ‘wfly-PCR-F1/R1’ (472bp) was divided into four overlapping regions of 315bp across a 50bp sliding window size, and the 518bp amplicon generated by the ‘wfly-PCR-F2/R2’ primer pair was divided into five overlapping regions of 315bp. We also assessed full amplicon lengths from both primer pairs (i.e., F1/R1: 472bp; F2/R2: 518bp, blue coloured cells). Cryptic species that could not be defined were represented by grey coloured cells, and sequences that were successfully defined into their respective species clades were shown by green cells. Dark grey cells indicated failed species delimitation by both primer pairs and full species delimitation will require the complete 657bp barcoding gene. The number of clean mtCOI sequences in the database (db) of Kunz, Tay, Elfekih, et al. (2019) for each species analysed are indicated (total: 224 sequences).

## Discussion

This study successfully demonstrated the applicability, transferability and powers of the multi-primer metabarcoding approach to investigate species compositions within an agroecological context, using the highly challenging *Bemisia* whitefly cryptic pest species as a study model system. These complementary sets of primers developed to amplify across the large genetic distances among the *B. tabaci* species complex also detected *non-Bemisia* hemipteran whitefly-nymphs, while showing potential to also simultaneously survey for associated parasitoid species diversity and parasitism rates, thereby further demonstrating the wide applicability and contribution to our understanding of trophic interactions between pests, beneficial insects and host plant usage across highly heterogeneous agricultural landscapes. Our multi-primer metabarcoding approach that involved high-volume sampled biological material demonstrated host plant preference and species composition differences in African *Bemisia* species (Table 3, Fig. 3), and that ascertaining a *Bemisia* species’ taxonomic status will require the standard length (i.e., ≥657bp) mtCOI barcoding 3’ gene region (Suppl. Table 1; Fig. 4), in addition to considering also developmental/life stages (Fig. 3F), and the species’ plant virus vectoring capabilities (e.g., Chi et al., 2020). We also outlined the development of an easy-to-use tailored bioinformatic analysis workflow, and provided the *Bemisia* research communities with two sets of highly efficient PCR primers that have the capability to amplify the alternative but widely used 3’ mtCOI *Bemisia* barcoding region. Our primer sets when used in metabarcoding of *Bemisia* nymphs, also detect various Aphelinid hymenopteran parasitoid species within the *Eretmocerus* and *Encarsia* genera. Unexpected co-amplification of various fungal and bacterial homologous gene regions, albeit at low frequencies, was also detected.

DNA metabarcoding approaches have been reported for integrative taxonomy (e.g., Cruaud, Rasplus, Rodriguez, & Cruaud, 2017; Sigut et al., 2017), their potential for invasive species surveys reviewed (Piper et al., 2019), for understanding relationships between species abundances and associated ecological factors (e.g., Macfadyen et al., 2021), and impact from aquacultural activities demonstrated (He et al., 2021). Adoption of such metabarcoding approaches for landscape ecological investigations under field conditions between insect pest hosts and beneficial insects may also be possible (e.g., Macfadyen et al., 2021), although appropriate experimental designs, primer improvement and modifications to analysis workflow including building of relevant barcoding database will be needed to increase species survey and detection sensitivity. For example, DNA barcoding primers for arthropods including the Hymenoptera (e.g., Folmer, Black, Hoeh, Lutz, & Vrijenhoek, 1994; Hebert, Penton, Burns, Janzen, & Hallwachs, 2004; see also Woodcock et al., 2013) could be included as internal checks to compare frequencies of parasitism and species diversity, with concurrent analysis following the HTS pipeline as outlined (Fig. 2). Studies that aimed to understand host-parasitoid interactions involving Aphalinid parasitoids faces other challenge such as high species diversity in the Aphelinidae (e.g., Gebiola et al., 2017; Hernández - Suárez et al., 2003), while different reproductive modes (sexual vs. asexual vs. Rickettsiales-induced sex ratio biases) in these haplodiploid parasitoids further complicate our understanding and interpretation of such biological interactions. Efficacies of these two complementary primer sets described in this study on hymenopteran parasitoids requires further assessment, as primer choice are known to affect species diversity recovery rates (e.g., Hajibabaei et al., 2019) and is currently beyond our research aims.

Primer efficacies is a significant issue for *Bemisia* molecular diagnostics (e.g., Elfekih, Tay, et al., 2018; Mugerwa et al., 2018; Shatters et al., 2009), and have contributed to mis-identification of pseudo-species (e.g., MEAM-K, Roopa et al., 2015; MEAM2, Delatte et al., 2005; Karut, Kaydan, Tok, Doker, & Kazak, 2015; Ueda, Kitamura, Kijima, Honda, & Kanmiya, 2009; SSA4, Berry et al., 2004) as demonstrated through genomic approaches (Elfekih et al., 2019; Kunz, Tay, Elfekih, et al., 2019; Tay, Elfekih, Court, et al., 2017; Vyskočilová et al., 2018). Our two primer sets while successful in test scenarios in amplifying diverse *Bemisia* cryptic species (Table 2), by themselves, have nevertheless failed to resolve species status within various cryptic species complexes (Fig. 4) due to insufficient DNA length. In a previous study by Wang et al. (2018), it was proposed that cost-effective NGS barcoding solutions were available albeit with shorter (e.g., 313bp) partial mtCOI sequences, however this would result in poor study outcome due to the biological and taxonomical complexity associated with the *B. tabaci* species complex (see Fig. 4). When used together however, was able to differentiate all but the closely related MED-ASL/MED-Burkina Faso species complex, SSA11, and also the two closely related NW2/NW2-2 species complex. Differentiating between certain cryptic *Bemisia* species may benefit from using alternative DNA markers and gene regions. For example, Vyskočilová et al. (2018) showed that species delimitation of the MED-ASL from other MED cryptic species was best achieved using the standard (i.e., 5’) barcoding gene region or via full mitogenomes. Furthermore, delimiting the true identity of the SSA1 species complex in our study sites will likely require genome-wide single nucleotide polymorphic markers (Elfekih et al., 2019). For the putative NW2/NW2-2 species complex identified by Kunz, Tay, Elfekih, et al. (2019), it remained to be seen if the 5’ COI gene region or if genome-wide single nucleotide polymorphic makers would prove to be the more effective diagnostic marker system.

Test scenarios involving evolutionarily divergent whitefly species (Tables 1 and 2) and field data on diversity of whitefly species detected (e.g., included novel *Bemisia* and *Trialeurodes* species; Table 3) suggested that overall non-detection rates of species could be potentially low. Our primers further demonstrated unexpected detection of novel *Encarsia* and *Eretmocerus* parasitoid species in the African cassava and non-cassava fields, however the true genetic and species diversity of these minute parasitoids will require independent assessment for confirmation, such as to include the standard barcode COI gene region using standard primers (e.g., C_LepFoIF/C_LepFoIR; Woodcock et al., 2013) as internal control. Nevertheless, understanding of trophic interactions between crop hosts, pest/beneficial insects and their potential biological control agents represents a molecular ecological network research area that could yield significant insights via metabarcoding approach (e.g., Bansch et al., 2020; Evans, Kitson, Lunt, Straw, & Pocock, 2016; Sow et al., 2019).

Determining which insect represents the major species within diverse agricultural landscape settings requires adequate population-wide sampling (Kalyebi et al., 2021; Kalyebi et al., 2018; Macfadyen et al., 2021). Conclusions based on few individuals sampled at a large number of sites are inherently problematic; there is a need, therefore, for the development of techniques to scale up data collection and processing of samples, and to develop associated bioinformatic workflows and analytical pipelines to provide robust findings at economic scales. For the cassava cultivation landscape in sub-Saharan Africa, determining the dominant pest whitefly cryptic species represents a significant challenge due to difficulties with species identification without the help of molecular DNA markers (De Barro et al., 2011; Kunz, Tay, Elfekih, et al., 2019). Furthermore, the often-high population sizes associated with individual fields, and poor PCR primer efficacies (Elfekih, Tay, et al., 2018; Mugerwa et al., 2018; Shatters et al., 2009) regularly resulted in suboptimal PCR amplification, and the difficulty encountered in sequence quality control (Kunz, Tay, Elfekih, et al., 2019; Tay, Elfekih, Court, et al., 2017; Vyskočilová et al., 2018), have been persistent problems in *Bemisia* molecular systematics and diagnostics.

In this study, we showed that by applying the DNA barcoding and the species ‘genetic gaps’ concepts first demonstrated by Dinsdale et al. (2010; see also Kunz, Tay, Elfekih, et al., 2019), and with careful re-designing of DNA markers for the HTS metabarcoding platform, it was possible to develop a highly efficient method for large-scale, effective and economic sampling and identification of the cryptic whitefly species to contribute to their management and control in challenging agricultural landscape settings (Kalyebi et al., 2018; Macfadyen et al., 2018; Macfadyen et al., 2021). Our species survey findings showed that SSA2 and *B. afer* were also present in the East Africa’s cassava landscape in addition to the dominant SSA1 species, and contrasted the reported absence of SSA2 based on sampling of low numbers of individuals in Ugandan cassava landscape (Ally et al., 2019). Co-exitance of both SSA1 and SSA2 may have important implication to genome evolution through introgression (Elfekih et al. 2021) and emergence of cassava disease epidemic (Legg, Sseruwagi, et al., 2014; Patil & Fauquet, 2009). Significant *Bemisia* whitefly species diversity such as the Asia species complex (e.g., see Kunz, Tay, Elfekih, et al., 2019) as well as invasive MED and MEAM1 species existed across the Asian continent. Adopting our metabarcoding approach to delimit species status, composition, prevalence, and their association with asymptomatic/diseased cassava plants could provide knowledge necessary to assist with the development of management strategies for the recent SLCMV disease outbreaks in the South East Asian region (Chi et al., 2020; Minato et al., 2019).

The importance of using standardised approach to sample sufficiently large quantities of *B. tabaci* cryptic species (Sseruwagi, Sserubombwe, Legg, Ndunguru, & Thresh, 2004; Macfadyen et al., 2021) especially across diverse agricultural landscape and between different life stages (e.g., nymph vs. adult) is also demonstrated (e.g., Table 3, Fig. 3F), where we detected significant differences in species compositions (e.g., between *B. afer* and SSA1). The approach developed here is therefore useful for processing nymphal data that may have strong links to host plants. Standardized sampling techniques for nymphs are less well-developed than for adult *B. tabaci* on cassava (which are considered easier to count and collect). In reality, the focus of the research question should drive the sampling approach selected (be that nymphs, parasitized nymphs, or adults); however, in all cases scaling up the numbers of individuals processed is critical. Another important distinction of specific sampling of whiteflies from different host crops was also demonstrated in cassava versus non-cassava host crops (Table 3, Figs. D, E), where differences in species compositions, especially between *B. afer* and SSA1 *cf*. MED, Ug, and IO, respectively, were evident. Ecological and biological factors therefore underpinning *Bemisia* species compositions in Africa agricultural landscapes. Similarly, ecological factors and agricultural practices will likely impact on pest species compositions in other agro-ecological landscape systems and can be assessed by adopting the multi-primer metabarcoding method outlined here.

While our method enables efficient ascertainment of species composition at the landscape level, this method should not be used to determine individual maternal lineages of mtDNA haplotypes due to the high sensitivity of detecting random PCR-introduced SNPs in individual amplicon molecules. At the landscape survey level, ascertaining specific maternal lineages does not represent a criterion needed to enable species identity to be determined, provided that sequence quality was adequately managed to avoid misidentification of species status due to mtDNA pseudogenes. Elfekih et al. (2021) have shown that NUMTs and pseudogenes could lead to significant misinterpretation of *Bemisia* population structure in the African cassava landscape regardless of whether partial mtCOI or genome-wide SNPs were used (see e.g., Wosula, Chen, Fei, & Legg, 2017). A significant limitation on our approach is the quality of the library used to interrogate the sequences generated. Sanger sequencing-generated partial mtCOI gene sequences in the public database (e.g., NCBI; see Vyskočilová et al. (2018) regarding MED-related pseudogenes/NUMTs; see also Kunz, Tay, Elfekih, et al. (2019) for reanalysis of *B. tabaci* DNA database (Boykin, Savill, & De Barro, 2017)) are gradually being recognised for containing high numbers of problematic sequences (see e.g., Nacer & do Amaral, 2017; Song, Buhay, Whiting, & Crandall, 2008; Triant & Dewoody, 2007). Such readily available public DNA global databases will need to be critically reviewed and assessed, given that a large scientific community relies on these sequences to assist with species identification.

A further complicating factor in molecular diagnostics of *Bemisia* species is the significant challenge associated with their nomenclature, cryptic taxonomy (Boykin et al., 2018; De Barro et al., 2011), and the preferred partial COI gene region widely adopted by the *Bemisia* tabaci research community (i.e., 3’ COI region) against the standard barcode 5’ COI region. For example, against the 34 *B. tabaci* cryptic species and five non-*tabaci* species presented by Kunz, Tay, Elfekih, et al. (2019) based on sanitized dataset, Barcode of Life data system v4 (BOLDSYSTEMS) did not identify 30 of these, while eight cryptic species were all identified as *B. tabaci*, and only one was correctly identified as *B. afer* (Suppl. Table 1). Furthermore, within the BOLDSYSTEMS database, the invasive MEAM1 was identified as *‘B. tabaci’*, while the Mediterranean (MED) species, shown as the real *B. tabaci* species as originally described by Gennadius in 1889 (Tay et al., 2012), returned as ‘no species level match’. While BOLDSYSTEMS database and the NCBI database are linked, inherent problems associated with reporting cryptic species remained (e.g., unable to provide cryptic species identity in NCBI sequence submission process) and will continue to impact on research outcomes especially as metabarcoding becomes more widely and rapidly adopted. Reporting of full COI gene such as through assemblies of draft mitogenomes in the *Bemisia* whitefly species complex (e.g., Kunz, Tay, Court, et al., 2019; Tay, Elfekih, Court, et al., 2017; Tay, Elfekih, Polaszek, et al., 2017; Thao & Baumann, 2004; Vyskočilová et al., 2018; Wang et al., 2016) can consolidate the different databases for *B. tabaci* cryptic species. Linking of other non-standard gene regions (e.g., mitochondrial DNA 12S rRNA and ND2 gene regions) to the standard 5’ COI gene region will also further resolve our understanding of species diversity from diverse ecosystems (e.g., Miya, Gotoh, & Sado, 2020; Stefanni et al., 2018; Weitemier et al., 2021).

Our metabarcoding platform can be adapted to understand other pest species complexes in agricultural systems and may be especially effective for invasive insect pests with small body size and where high density of population sizes can build up rapidly, such as in invasive whiteflies, thrips, mites, and aphids. We propose, therefore, to apply this approach to: (1) greatly increase number of individuals that can be sampled, processed and characterised simultaneously for their widely applied proposed 5’ COI ( Hebert et al., 2003) or alternative ‘DNA barcoding’ genes and gene regions (e.g., the mtCOI 3’ region), and the earlier (Woese, Kandler, & Wheelis, 1990) proposed RNA genes and related regions (i.e., ITS rRNA region for fungi, e.g., Xu (2016); Microsporidia, e.g., Ghosh & Weiss, 2009; Klee, Tay, & Paxton, 2006; O’Mahony, Tay, & Paxton, 2007; Tay, O’Mahony, & Paxton, 2005; Velasquez et al., 1996; 16S rRNA, e.g., Shelomi & Chen, 2020; 18S rRNA, e.g., Popovic et al., 2018; 28S rRNA, e.g., Pawlowski et al., 2012); (2) estimate individual species proportion across large agricultural landscapes, and (3) in nymphal/larval samples, identify parasitoid species that target the agricultural pests of interest. Contrasting the majority of metabarcoding studies published to-date, our approach utilises significantly longer partial COI gene sequence through two sets of complementary primer sets (i.e., *ca*. 650bp *cf*. 130-421bp; Brandon-Mong et al., 2015; Hajibabaei et al., 2019; Miya et al., 2020; Stefanni et al., 2018) which was important for defining cryptic *Bemisia* species status. The use of this multi-primer metabarcoding approach to generate longer sequence length and targeting the 5’ COI barcoding gene region could enable greater species identification with the support of the BOLDSYSTEMS.

While the high-throughput amplicon sequencing method in a species composition study in agroecological landscapes represents a relatively new field of research, the use of this method to identify genes of biosecurity importance has been demonstrated for understanding the spread of pyrethroid resistance genes in the red-legged earth mites *Halotydeus destructor* (Edwards et al., 2018). The method described by Edwards et al. (2018) can be incorporated into our high-throughput amplicon sequencing platform for rapid species composition identification and insecticide resistance allele frequency estimates. This would enable a genomic approach that would simultaneously allow the identification of species composition at landscape scales, the survey of insecticide resistance profiles (e.g., Hopkinson et al., 2019), and assessment of the implication of chemical control strategies on diversity and density of beneficial insects such as parasitoids. Such study would be especially informative when temporal scales are included within the study designs (e.g., Kalyebi et al., 2021; Macfadyen et al., 2021).

Developmental stages sampled do impact *Bemisia* species compositions and the conclusions drawn from a survey, as demonstrated in the Malawi cassava sampling sites for adult and nymphal *Bemisia* populations. The extent of such variability in target species composition between immature and adult life stages for many other biological systems can also benefit from the HTS metabarcoding method described here. In this study, *Bemisia* nymphs that visibly showed signs of being parasitised by hymenopteran parasitoids were first excluded prior to genomic processing. Despite this, *Encarsia* and *Eretmocerus* parasitoid mtCOI sequences were detected, thereby highlighting the importance of biological control of *Bemisia* pest species by parasitoids in the sub-Saharan African agricultural system. While we have not optimised this workflow to address host-parasitoid interaction research questions, we have demonstrated the power of HTS genomic approaches to document and facilitate an understanding of these biological interactions. Improved understanding of the ecological impact of pest species systems from genomic perspectives will further transform our understanding of sustainable agricultural production. Appropriate experimental designs including careful development of sampling procedures will be needed for meaningful interpretation that takes advantage of the large volume of genomic data generated by the HTS platform. Our study demonstrates the successful use of multi-primer HTS metabarcoding methods in classical field-ecological studies and for novel cryptic species discovery. The adoption of this approach more widely has the potential to help integrate global biodiversity with genomic (Arribas, Andujar, Bidartondo, & al., 2021) and transform global biosecurity preparedness to impact understanding of ecological network, conservation biology, human health, and food security.

## Acknowledgements

Laboratory colonies of whitefly samples (*Bemisia* SSA1, SSA2, SSA3) were provided by the NRI, University of Greenwich, UK. J. Ryu (CSIRO) assisted with sample sorting, C. Paull and A. Hulthen (CSIRO) assisted with field surveys of African cassava whitefly samples. We thank Dr Andy Polaszek (NHM, UK) for discussion on the *Encarsisa* and *Eretmocerus* parasitoids detected in the *Bemisia* hosts. This work was supported by the Natural Resources Institute, University of Greenwich from a grant provided by the Bill & Melinda Gates foundation (Grant Agreement OPP1058938). We would like to thank the broader African cassava whitefly project team for their enthusiasm and support throughout this study.

## Data Availability Statement

The data that support the findings of this study are available from GenBank (GenBank accession numbers cited throughout). Raw HTS sequence data can be downloaded from GenBank (SRA xxxx01, xxxx02, xxxx03).

