## Supplementary material for "A high-throughput amplicon sequencing approach for population-wide species diversity and composition survey": Suppl. Material I

This method of screening for *Bemisia* species complex mitochondrial DNA cytochrome oxidase subunit I (mtDNA COI) haplotype diversity at the population level aims to utilise the high-throughput sequencing (HTS) capacity to rapidly estimate proportions of haplotype diversity in set numbers of individuals sampled from agricultural landscapes. The protocol has the greatest efficacies when target pest organisms sampled were stored immediately in >95% ethanol and kept at -18˚C to -20˚C prior to sorting of nymphs/adult whiteflies for gDNA extraction. We used the Illumina MiSeq sequencing platform to process all amplicon libraries.

For all steps, use sterile techniques to avoid cross-contamination.

**I.	Field samples and genomic (gDNA) extraction**

1.	For each population: pool between 20 or 40 *Bemisia* adults/nymphs into a 1.5 mL sterile Eppendorf tubes, top up (500 µL–1000 µL) with >95% ethanol and store in -18˚C until nymphs from multiple populations have been pooled. (**Note:** record adult/nymph sample codes used in gDNA extraction for each population).

2.	Once sufficient numbers of whiteflies have been sorted from desired number of populations (i.e., having >48 tubes representing 48 individual populations, and each tube containing 20–40 nymphs), discard the ethanol from the tube and briefly dry (5–10 min) at 56˚C to remove all traces of the ethanol.

3.	Extract gDNA from pooled samples using Qiagen Blood & Tissue DNA extraction (BTDE) kit (Cat. # 69506) following supplier’s recommended protocol. Allow 24–48hrs for initial gDNA digestion, and remove RNA using recommended amount of RNase A (Qiagen Cat No. 19101) as per Qiagen BTDE kit protocol.

4.	For each pooled population, elute gDNA from the Qiagen BTDE kit using 200 µL of Buffer EB Elution buffer (Qiagen Cat. No. 19086) to minimise buffer incompatibility with the downstream NGS protocol.

5.	Quantify gDNA concentration for each pooled sample using 2 µL of the eluted gDNA in a Qubit fluorometer (ThermoFisher Scientific).

6.	Standardise gDNA concentration per population to 5 ng/µL using Buffer EB.

7.	Store standardised gDNA (5 ng/µL) at -20˚C (in a freezer) until ready for PCR amplicon library preparation.

**II.	PCR amplicon DNA library preparation for ILLUMINA MiSeq HTS platform**

This step takes the pooled gDNA from each population and amplifies the mtDNA COI partial gene for identification of *Bemisia* cryptic species complex, and to assign haplotypes and species complex based on the mtCOI database of Kunz et al. (2019). The Illumina NGS protocol for **16S Metagenomic Sequencing Library Preparation** ([Part # 15044223 Rev. B](http://www.illumina.com/content/dam/illumina-support/documents/documentation/chemistry_documentation/16s/16s-metagenomic-library-prep-guide-15044223-b.pdf)) is adapted to enable HTS amplicon sequencing of *Bemisia* mtDNA COI partial gene. Where modifications have been made to Illumina’s **16S Metagenomic Sequencing Library Preparation** protocol, these are specifically detailed below.

HTS PCR-amplicon sequencing of the 657bp *Bemisia* mtDNA COI region involves preparation of two amplicon libraries (consisting of amplicons from primers F1/R1, and F2/R2; **Fig. 1**) for each population. This two-steps PCR involving two sets of overlapping primers (Table 1) serves as a ‘back-up’ system to minimise the likelihood of complete PCR failure of a particular population sample, such as due to difficulties encountered with primer efficacies especially for *Bemisia* whiteflies, enables the complete 657bp region to be sequenced, and trimming of poor quality amplicon regions introduced during the DNA template extension stage by DNA polymerases.

**Suppl. Fig. 1.** Schematic representation of the 1^st^ round PCR products generated during the construction of NGS amplicon libraries for *Bemisia* whitefly partial (657 bp) mtDNA COI barcode region. For each population of *Bemisia* whitefly, two amplicon libraries (F1/R1 and F2/R2) are constructed in 2 successive rounds of PCR so as to cover the targeted 657 bp region; The 1^st^ round of PCR generates F1/R1 and F2/F2 COI PCR products each with Illumina overhang sequences (COI primer sequences shown as red arrows, Illumina overhang sequences as grey tails). The F1/R1 overhang amplicons are 645 bp (i.e., 578 bp mtCOI + 33bp F1 adaptor sequence + 34 bp R1 adaptor sequence). The F2/R2 overhang amplicons are 663 bp (i.e., 596 bp mtCOI + 33 bp F2 adaptor sequence + 34 bp R2 adaptor sequence); The 2^nd^ round of PCR then uses the 1^st^ round amplicons as template and targets their 5’ Illumina overhang sequences with commercially available Illumina Nextera-XT indexing primers. This limited cycle PCR step adds on the required Illumina sequencing adapters and indexing barcode sequences and completes the construction process for the F1/R1 and F2/R2 amplicon libraries.

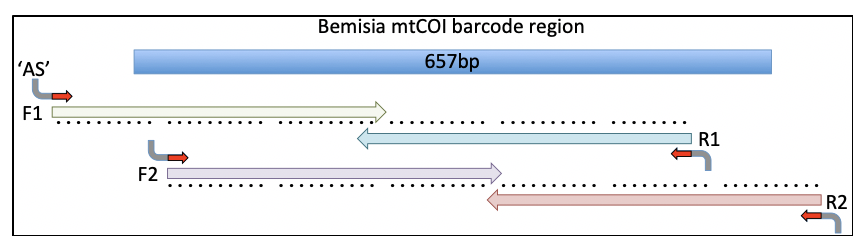

**Table 1:** Primer sequences and expected amplicon size (in base pairs, ‘bp’) for generating 1^st^ round PCR products for the construction of HTS amplicon libraries. The 5’ Illumina overhang sequences are not underlined, the *Bemisia* specific primer sequences are underlined. Ambiguous bases are R (G/A), W(T/A), Y(C/T).

| **Primer** | **Primer sequences** | **Amplicon** |
| --- | --- | --- |
| wfly-PCR-F1 | TCGTCGGCAGCGTCAGATGTGTATAAGAGACAGTGGTTYTTTGGTCATCCRGAAG | 645 bp |
| wfly-PCR-R1 | GTCTCGTGGGCTCGGAGATGTGTATAAGAGACAGGGAAARAAWGTTAARTTWACTCC |  |
| wfly-PCR-F2 | TCGTCGGCAGCGTCAGATGTGTATAAGAGACAGCGRGCTTAYTTYACTTCAGCYAC | 663 bp |
| wfly-PCR-R2 | GTCTCGTGGGCTCGGAGATGTGTATAAGAGACAGGGYTTATTRATTTTYCAYTCTA |  |

1.	For each pooled gDNA sample, two separate 1^st^ round PCR amplifications using (i) wfly-PCR-F1/R1, and (ii) wfly-PCR-F2/R2 were carried out. PCR procedures follow ‘Amplicon PCR’ step (**Pages 6-7**) of the Illumina **16S Metagenomic Sequence Library Preparation** protocol with slight modifications as detailed below. PCR profiles for both F1/R1 and F2/R2 are identical, and consisted of:

• 95˚C for 5 minutes

• 30cycles of

• 95˚C for 30 seconds

• 48˚C for 30 seconds (optimised for primers in Table 1)

• 72˚C for 30 seconds

• 72˚C for 5 minutes

• Hold at 4˚C

2.	Determine the success of PCR amplification for all samples by gel electrophoresis, using 5µL of PCR amplicon mixed with 1µL 6x loading dye and loaded on a 1.25% agarose gel.

3.	Perform PCR Clean-Up by following the Illumina **16S Metagenomic Sequence Library Preparation** protocol (**Pages 8**–**9**), ensuring that correct AMPure XP bead ratios are used for the clean-up. For the current protocol, our experience indicated a bead ratio of 0.8 is sufficient for the PCR clean up, although this would likely vary between laboratories and PCR polymerase used.

4.	Quantify PCR amplicon concentration using 2 µL of amplicon on a Qubit fluorometer. This step determines if sufficient amplicons were generated for subsequent PCR clean-up and Index-PCR steps.

5.	Ascertain amplicon sizes on an Agilent Technologies TapeStation using the High Sensitivity D1000 ScreenTape, following instructions as recommended by the manufacturer. Amplicon sizes should match the expected amplicon sizes listed in Table 1.

6.	Follow through to step 16 of the PCR Clean-Up protocol (see Illumina **16S Metagenomic Sequence Library Preparation** protocol, **Page 9**)L.

7.	Index PCR (**Pages 10**–**12**): As per Illumina’s **16S Metagenomic Sequence Library Preparation** protocol.

8.	PCR Clean-up 2 (**Pages 13**–**15**):

[i]	After the Index PCR and PCR clean-up steps, using a Qubit flurometer to quantify the post purification amplicon concentration (use 2 µL of amplicon).

[ii]	Standardise Index PCR amplicon concentrations for all samples as required

[iii]	Combine equal amount (e.g., 100 ng/sample) of F1/R1 index PCR amplicon products into a single Eppendorf tube.

[iv]	Combine equal amount (e.g., 100 ng/sample) of F2/R2 index PCR amplicon products into a single Eppendorf tube.

[v]	Determine the amplicon sizes of both pooled F1/R1 and F2/R2 index PCR amplicon products on an Agilent Technologies TapeStation. Include a sample each of F1/R1 and F2/R2 pre-Index PCR amplicon templates as controls to help ascertain if index PCR had been successful (**Note:** there should be a noticeable increase (i.e., by ~69 bp) in amplicon sizes for Index PCR amplicons as compare with pre-index PCR amplicon).

[vi]	Run pooled index-PCR on a 1.25% low melt agarose gel, excise expected amplicon fragments and perform a gel clean-up using commercial gel purification kit (e.g., the Zymoclean Gel DNA Recovery Kit), and elute purified amplicon in 22.5µL EB.

[vii]	Repeat TapeStation analysis to ensure the gel purification step has successfully removed all smaller fragments of PCR duplexes.

[viii]	Normalise purified index PCR amplicon to 4 nM. The PCR amplicon products are now ready for the MiSeq run (on a MiSeq Reagent kits version 3).
