## Supplementary material for "A high-throughput amplicon sequencing approach for population-wide species diversity and composition survey": Suppl. Table 1

**Suppl. Table 1:** Selected GenBank accession numbers representing the current 34 known species of *Bemisia tabaci* cryptic species (#1-#34) as well five selected non-*tabaci* species (#34-#39) as reported by {Kunz, 2019 #23}. MEAM1 (#20) was identified as ‘*B. tabaci*’ within the Barcode of Life Data Systems (BOLDSYSTEMS) database, however the true *B. tabaci* (#23 ‘MED’; {Tay, 2012 #26} was instead not identified and reported as “a species level match could not be made. The nearest match is *Bemisia tabaci*”, showing that molecular diagnostics of *B. tabaci* cryptic species complex based on the BOLDSYSTEMS database remained challenging and that primer validation was likewise not possible. † BOLDSYSTEMS record from GenBank full mtCOI gene sequence (• 1537-1539bp; • 725-900bp). Similarity: no percentage results provided based on Species Level Barcode Records (i.e., COI Species Database), per centage provided based on COI Full Database. BOLDSYSTEMS Best ID identified as Bemisia tabaci based on Species Level Barcode Records, when ‘No Match’ was identified, top match was determined based on COI Full Database.

|  | **GenBank** | **Species** | **BOLD Best ID** | **Similarity** | **Comment** | **BOLD Sequence ID** | **†** |
| --- | --- | --- | --- | --- | --- | --- | --- |
| **1** | JF901839 | NewWorld2-2 | *Bemisia tabaci* | 98.99% | No species level match; Nearest: *B. tabaci* (LR535718) | GBMNA33230-19 | • |
| **2** | EU427729 | New World 1 | No Match | 95.96% | Top match: *B. tabaci*, NC_006279, Mined from GenBank, NCBI. Locus: NADH dehydrogenase subunit 4L (282bp) | GBMTG494-16 |  |
| **3** | AJ748398 | Asia 1 | *Bemisia tabaci* | 98.99% | No species level match; Nearest: *B. tabaci* (MK490832) | GBMNA45551-19 | • |
| **4** | KC113546 | China2 | No Match | 91.67% | Top match: *B. tabaci*, Mined from GenBank, NCBI (FJ939624) | GBMIN9096-12.COI-5P | • |
| **5** | GQ139495 | China1 | No Match | 91.67% | Top match: *B. tabaci*, Mined from GenBank, NCBI (FJ939624) | GBMIN9096-12.COI-5P | • |
| **6** | GU086328 | Australia & Australia II | *Bemisia tabaci* | 98.99% | No species level match; Nearest: *B. tabaci* (KY951451) | GBMNA17955-19 | • |
| **7** | HM137337 | AsiaII_1 | *Bemisia tabaci* | 98.96% | No species level match; Nearest: *B. tabaci* (KX810024) | GBMHH13664-19 | • |
| **8** | AJ783706 | AsiaII_3 | No Match | 90.63% | Top match: *B. tabaci*, Mined from GenBank, NCBI (KX810024) | GBMHH13664-19 | • |
| **9** | AJ748391 | AsiaII_5 | No Match | 88.89% | Top match: *B. tabaci*, Mined from GenBank, NCBI (KX810024) | GBMHH13664-19 | • |
| **10** | DQ174520 | AsiaII_6 | No Match | 90.91% | Top match: *B. tabaci*, Mined from GenBank, NCBI (KX810024) | GBMHH13664-19 | • |
| **11** | GQ139492 | *Bemisia emiliae*/AsiaII_7 | No Match | 90.91% | Top match: *B. tabaci*, Mined from GenBank, NCBI (KX810024) | GBMHH13664-19 | • |
| **12** | HM590188 | AsiaII_8 | No Match | 91.92% | Top match: *Bemisia sp. WTT-2017*, Mined from GenBank, NCBI (KX714968) | GBMNA17746-19 | • |
| **13** | HM137313 | AsiaII_9 | No Match | 91.67% | Top match: *B. tabaci*, NC_006279, Mined from GenBank, NCBI. Locus: NADH dehydrogenase subunit 4L (282bp) | GBMTG491-16 |  |
| **14** | HM137356 | AsiaII_10 | No Match | 87.88% | Top match: *B. tabaci*, Mined from GenBank, NCBI (KX810024) | GBMHH13664-19 | • |
| **15** | HM590147 | AsiaII_11 | No Match | 89.9% | Top match: *Bemisia sp. WTT-2017*, Mined from GenBank, NCBI (KX714968) | GBMNA17746-19 | • |
| **16** | AY827601 | Italy1 | No Match | 88.89% | Top match: *B. tabaci*, Mined from GenBank, NCBI (MK490832) | GBMNA45551-19 | • |
| **17** | AY057180 | Sub-Saharan 1 | *Bemisia tabaci* | 98.96% | No species level ID; Nearest: *B. tabaci* (LR535717) | GBMNA33229-19 | • |
| **18** | AY057173 | Sub-Saharan 2 | No Match | 91.92% | No species level ID; Nearest: *B. tabaci* (LR535717) | GBMNA33229-19 | • |
| **19** | KM377923 | Sub-Saharan 3 | No Match | 91.92% | No species level ID; Nearest: *B. tabaci* (LR535717) | GBMNA33229-19 | • |
| **20** | HM070414 | MEAM1 | *Bemisia tabaci* | 100% | Matched to *Bemisia tabaci* (FJ939628) | GBMIN9094-12.COI-5P | • |
| **21** | AJ550171 | Indian Ocean | *Bemisia tabaci* | 98.99% | No species level match; Nearest: *B. tabaci* (KY951448) | GBMNA17952-19 | • |
| **22** | AY903576 | Uganda | No Match | 84.38% | Top match: *B. tabaci*, Mined from GenBank, NCBI (FJ939628) | GBMIN9094-12 | • |
| **23** | FJ766381 | MED | *Bemisia tabaci* | 98.96% | No species level match; Nearest: *B. tabaci* (FJ939624) | GBMIN9096-12 | • |
| **24** | DQ174527 | Asia_III | No Match | 92.71% | No species level match; Nearest: *B. tabaci* (MK490832) | GBMNA45551-19 | • |
| **25** | EU760739 | Africa_Cameroon | No Match | 91.11% | No species level match; Nearest: *B. tabaci* (KY951451) | GBMNA17955-19 | • |
| **26** | AB308111 | Japan2 ‘JpL’ | No Match | 96.97% | No species level match; Nearest: *B.* sp. WTT-2017 (KX714968) | GBMNA17746-19 | • |
| **27** | AB440786 | Japan1 | No Match | 92.71% | Top match: *B. tabaci*, Mined from GenBank, NCBI (MK490832) | GBMNA45551-19 | • |
| **28** | KX570778 | African_MEAM2 | No Match | 96.88% | Top match: *B. tabaci*, Mined from GenBank, NCBI (FJ939630) | GBMIN9093-12 | • |
| **29** | KX570804 | Sub-Saharan 10 | No Match | 89.58% | Top match: *B. tabaci*, Mined from GenBank, NCBI (FJ939629) | GBMIN9072-12 | • |
| **30** | KX570818 | Sub-Saharan 12 | No Match | 93.33% | Top match: *B. tabaci*, Mined from GenBank, NCBI (MH205754) | GBMHH14682-19 | • |
| **31** | KX570829 | Sub-Saharan 13 | No Match | 95.83% | Top match: *B. tabaci*, Mined from GenBank, NCBI (FJ939628) | GBMIN9094-12 | • |
| **32** | KX570852 | Sub-Saharan 6 | No Match | 95.83% | Top match: *B. tabaci*, Mined from GenBank, NCBI (LR535717) | GBMNA33229-19 | • |
| **33** | KX570856 | Sub-Saharan 9 | No Match | 93.94% | Top match: *B. tabaci*, Mined from GenBank, NCBI (LR535717) | GBMNA33229-19 | • |
| **34** | KX570815 | Sub-Saharan 11 | No Match | 88.89% | Top match: *B. tabaci*, Mined from GenBank, NCBI (KU877168) | GBMHH13663-19 | • |
| **35** | KF734668 | Bemisia afer | *Bemisia afer* | 98.99% | No species level match; Nearest: *B. afer* (NC_024056) Mined from GenBank, NCBI. Locus: NADH dehydrogenase subunit 4L (282bp) | GBMTG4823-16 |  |
| **36** | GU086362 | Bemisia atriplex | No Match | 89.58% | Top match: *B. tabaci*, Mined from GenBank, NCBI (MK490832) | GBMNA45551-19 | • |
| **37** | HQ457046 | Bemisia berbericola | No Match | 84.85% | Top match: *Conus* *ermineus* Mined from GenBank, NCBI (MH204531). Note: Gastropod | GBML16828-19 | • |
| **38** | MN056066 | Bemisia_sp_PDB_1 | No Match | 93.75% | Top match: *Bemisia sp. WTT-2017*, Mined from GenBank, NCBI (KX714968) | GBMNA17746-19 | • |
| **39** | MN056067 | Bemisia_sp_PDB_2-1 | No Match | 85.42% | Top match: *Aleurochiton aceris*, Mined from GenBank, NCBI (NC_006160) Locus: NADH dehydrogenase subunit 4L (282bp) | GBMTG491-16 |  |

**Note:** BOLDSYSTEMS search result where a species level match could not be made is indicated by ‘?’. Nearest species match is determined by the closest matching BIN of within 3%.

**References**
